# Bone Mineralization Regulation: Using Zebrafish as a Model to Study ANKH-associated Mineralization Disorders

**DOI:** 10.1101/2024.03.21.586098

**Authors:** Nuwanthika Wathuliyadde, Katherine E. Willmore, Gregory M. Kelly

**Affiliations:** Department of Biology, Western University, London, Canada; Department of Anatomy and Cell Biology, Western University, London, Canada

**Keywords:** *ANKH* genes, phylogeny, paralog, gene expression, Bone mineralization; Craniometaphyseal Dysplasia, Zebrafish

## Abstract

Craniometaphyseal Dysplasia (CMD) is a rare skeletal disorder that can result from mutations in the *ANKH* gene. This gene encodes progressive ankylosis (ANK), which is responsible for transporting inorganic pyrophosphate (PPi) and ATP from the intracellular to the extracellular environment, where PPi inhibits bone mineralization. When ANK is dysfunctional, as in patients with CMD, the passage of PPi to the extracellular environment is reduced, leading to excess mineralization, particularly in bones of the skull. Zebrafish may serve as a promising model to study the mechanistic basis of CMD. Here we provide a detailed analysis of the zebrafish ankh paralogs, ankha and ankhb, in terms of their phylogenic relationship with ANKH in other vertebrates as well as their spatiotemporal expression patterns during zebrafish development. We found a closer evolutionary relationship exists between the zebrafish ankhb protein and its human and other “higher” vertebrate counterparts. Furthermore, we noted distinct temporal expression patterns with *ankha* more prominently expressed in early development stages, and *ankhb* expression at larval growth stages. Whole mount *in situ* hybridization was used to compare spatial expression patterns of each paralog during bone development. Both paralogs showed strong expression in the craniofacial region as well as the notochord and somites, with only subtle patterning differences. Given the substantial overlap in spatiotemporal expression of *ankha* and *ankhb*, the exact roles of these genes remain speculative. However, this study lays the groundwork for functional analyses of each ankh paralog and the potential of using zebrafish to find possible targeted therapies for CMD.

## 1. Introduction

Craniometaphyseal Dysplasia (CMD) is a rare disorder characterized by excessive mineralization of the skull and long bones [1]. Clinical features of this disease include flared long bone metaphyses, and several skull anomalies such as a wide nasal bridge, paranasal bossing, increased bizygomatic width and a prominent mandible [2]. Though less obvious phenotypically, the most severe functional consequences of CMD arise from progressive thickening of the cranial base. This excess mineralization can occlude the foramina through which the cranial nerves pass and lead to facial palsy, hearing loss and blindness [3]. Unfortunately, there is no permanent cure for CMD, and current treatment of the disease involves repetitive surgical recontouring of the affected bones [4]. Thus, further research is required to understand the mechanistic basis of CMD and to uncover potential therapeutic targets that can be used to prevent or permanently treat this disorder.

Genetic screens of individuals with CMD have determined that the disease can be caused by autosomal recessive mutations in the gene encoding the gap junctional protein connexin 43, but most commonly it is caused by autosomal dominant mutations in the gene progressive ankylosis homolog (*ANKH*) [5]. *ANKH* encodes a twelve pass transmembrane protein (ANK) and is expressed in tissues throughout the body including skeletal tissues, consistent with the bony phenotypes present in patients with CMD [6]. The function of ANK is to transport intracellular ATP and inorganic pyrophosphate (PPi) into the extracellular environment [7]. Within the extracellular environment, ATP can be hydrolyzed into AMP and PPi by ectonucleotide pyrophosphatase 1, further increasing the level of extracellular PPi (ePPi) [8]. ePPi acts as a potent inhibitor of bone mineralization, with low concentrations leading to excessive deposition of hydroxyapatite crystals [9,10]. Thus, through its role in transporting ATP and PPi to the extracellular environment, ANK helps to regulate ePPi concentration and ultimately bone mineralization [4, 8-10]. In CMD patients with *ANKH* mutations, the transport function of ANK is disrupted, causing excessive mineralization [1]. Several *ANKH* mutations have been reported in the CMD population, but most involve single amino acid substitutions, insertions or deletions and are located in the highly conserved cytosolic domain [6]. The most common mutation among patients with CMD involves the deletion of phenylalanine (position 377) of the polypeptide [4,6,11]. While patient screening has deepened our understanding of the genetic basis of CMD, animal models enable researchers to determine the cellular and molecular processes that underlie this disease and provide a means for exploring potential therapeutic targets.

Mouse models have greatly improved our understanding of the role of Ank *in vivo*. Loss of function mouse models of Ank (Ank^ank/ank^ and Ank^null/null^), demonstrate skeletal abnormalities including progressive arthritis and joint fusion that have been attributed to disrupted PPi transport and increased hydroxyapatite deposition within the joints [12,13]. The knockout mouse model of Ank^null/null^, mimics several characteristics observed in human CMD patients, such as, thickened skull bones, a narrowed foramen magnum and fused ossicles of the middle ear [13]. However, some CMD phenotypes are not present in these mice, including overgrown mandibles, obstructed nasal sinuses and flared long bone metaphyses [13]. Thus, Chen and colleagues developed a more targeted knock-in mouse model (Ank^KI/KI^) using an in-frame deletion of Phe377. Indeed, these mice closely phenocopy the symptoms observed in human CMD, including thick skull bones, a narrowed foramen magnum and fused middle ear bones, but additionally they demonstrate obliterated nasal sinuses, stenosis of cranial nerve foramina, widened long bone metaphyses and mandible hyperostosis [4]. While these mouse models have proven effective for helping to uncover the role of Ank and the pathogenesis of CMD, they are not perfect surrogates for the human disease. For example, in humans with CMD caused by mutations in *ANKH*, the disease arises in an autosomal dominant manner [6]. However, Ank^KI/+^ mice have skeletal phenotypes intermediate to wildtype (Ank^+/+^) and homozygous mutants (Ank^KI/KI^), with only Ank^KI/KI^ mutants demonstrating the phenotypic severity observed in patients suffering from CMD [4]. Additionally, joint stiffness observed in these mice has not been reported in CMD patients, indicating potential differences in disease manifestation between the mouse model and human CMD [4]. Moreover, mouse models are not as conducive to high-throughput therapeutic screens or studies of embryological pathogenesis of disease as other non-mammalian animal models. One such animal model commonly used to study bone diseases is the zebrafish [14].

Zebrafish offer convenience and relative cost-effectiveness compared to rodent models. For example, zebrafish produce a large clutch of embryos that are relatively transparent, develop rapidly and can be easily manipulated. Additionally, the zebrafish genome is sequenced and annotated and can be manipulated using knockout, knockdown and knock-in models of genes of interest to generate targeted tissue dysfunction [17-21]. Zebrafish have also proven to be an effective model organism for bone disease studies, with bone mineralization beginning five days post-fertilization (dpf) [14]. Thus, the utility of the zebrafish is well-known and could be exploited as a useful tool for studying the pathogenesis of CMD and targeted drug screening for this disease. However, zebrafish possess two paralogous *ankh* genes, *ankha* and *ankhb*, that emerged during a duplication event [10,19]. Given this gene duplication, a better understanding of the origin and evolutionary relationship of the *ankha* and *ankhb* paralogous with *Ankh/ANKH* of higher vertebrates, as well as their spatiotemporal expression patterns, is crucial to determine the efficacy of using zebrafish to model CMD.

While zebrafish are acknowledged as an outstanding model organism, a detailed understanding of the *ankh* genes within this species has yet to be established. Therefore, the purpose of our study was to first examine the evolutionary lineage of these paralogous genes present in zebrafish and assess their similarity to the human and rodent genes. Conducting a detailed investigation of *ankha* and *ankhb* in zebrafish using RT-PCR and whole mount *in situ* hybridization to understand their temporal and spatial expression revealed overlapping patterns, which complicated matters. Nevertheless, future research seeks to bridge this gap, enhancing our comprehension of the *ankh* genes in zebrafish and provide a rationale for using this model to study a rare human disease addressing ankh-associated mineralization disorders. Towards that end, we will be using CRISPR-Cas9 gene editing tools to knockout the *ankh* genes in zebrafish as we feel this approach will provide insights that address the inconsistencies seen in mice. Ultimately, our goal is to lead to a better understanding of how CMD can be alleviated.

## 2. Materials and Methods

### 2.1 Sequence Alignment and Phylogenetic Tree Construction of ANKH Proteins from Different Vertebrate Model Species

Phylogenetic trees were constructed to explore the evolutionary conservation and diversification of ANKH across vertebrates, from jawless fish to mammals, by obtaining and comparing their amino acid sequences from Ensembl [20]. The following sequences were used: Human (*Homo sapiens* - ENSP00000284268.6), Mouse (*Mus musculus* - ENSMUSP00000022875.7), Rat (*Rattus norvegicus* - ENSRNOP00055036996.1), Guinea Pig (*Cavia porcellus* - ENSCPOP00000011102.2), Rhesus macaque (*Macaca mulatta* - ENSMMUP00000080342.1), Pig (*Sus scrofa* - ENSSSCP00000017779.3), Domestic chicken (*Gallus gallus* – ENSGALT00010001547.1), Frog (*Xenopus tropicalis* - ENSXETP00000100546.1), Zebrafish (*Danio rerio* - ENSDARP00000105177.1 and ENSDARP00000002526.7), Common carp (*Cyprinus carpio -* ENSCCRT00000085173.2 and ENSCCRT00000029117.2), Mexican tetra (*Astyanax mexicanus ENSAMXG00000002808 and ENSAMXT00000038996.1), Hagfish (Myxini-ENSEBUG00000011035), Lamprey (Petromyzon marinus –ENSPMAT00000002489.1), and Elephant Shark (Australian Ghostshark) (Callorhinchus milii – ENSCMIT00000028310.1)*. The phylogenetic trees were constructed using neighbor-joining and maximum likelihood analyses to assess the taxonomic relationships of ANKH. The tree (Figure 1a) was rooted with *Hagfish (primitive species)* and was based on a multiple sequence alignment using MEGA version 10. The analysis was conducted with 10,000 bootstrap replicates [21].

**Figure 1.**
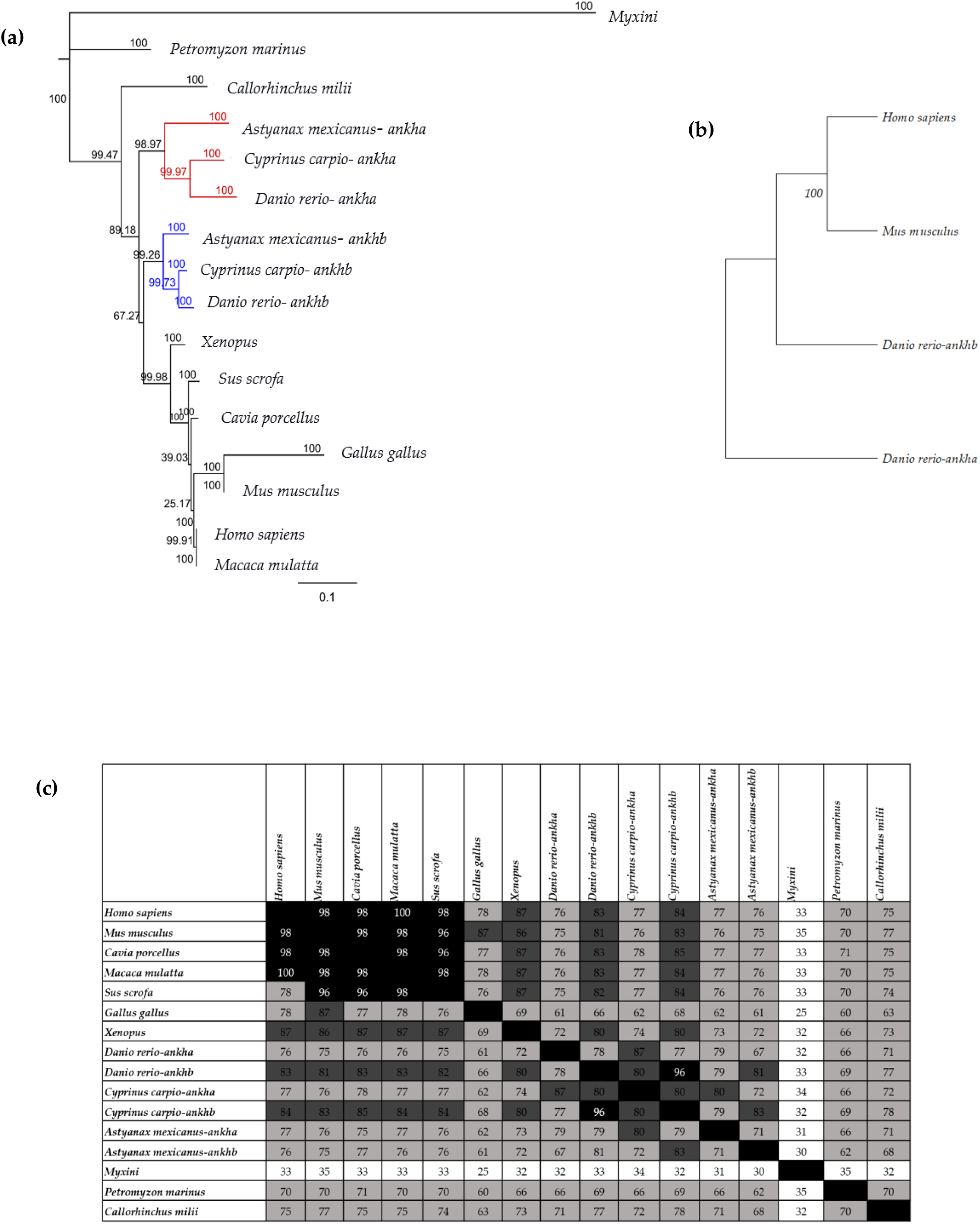
Phylogenetic Relationships of ANKH amino acid sequence in Vertebrate Model Organisms. (a) Results from neighbor-joining tree analysis using the out-group taxa (*Myxini*) demonstrates the relationships between vertebrate model organisms. Bootstrap analysis was performed using 10,000 replicates and these bootstrap values are included. Monophyletic clustering of teleost fish for ankha and ankhb are represented by red and blue branches, respectively. (b) The maximum likelihood tree includes human (*Homo sapien*), mouse (*Mus musculus*), and zebrafish (*Danio rerio*) *ankha* and *ankhb* and includes values from 10,000 replicate bootstrap analysis. (*c*) Matrix of pairwise amino acid sequence identities for ankh/ankha and ankhb across various vertebrate species determined using percentage identity matrix analysis. Boxes with a black background that have no values represent self-comparisons. High sequence identity is represented by darker shades whereas lighter shades highlight comparisons with lower sequence identity. Squares that have 90 to 100% identity are shown with a black background, sequence identity values from 80-90% are represented in dark gray, sequence identity values from 50-80% in light gray, and sequence identity values below 50% are represented with a white background. Each identity value is rounded to the nearest whole number.

### 2.2 Percentage identity matrix

The percentage identity matrix quantifies and visually presents the sequence identity between the ANKH amino acid sequences across different vertebrate species. This matrix was created by analyzing amino acid sequences specified in Section 2.1. The amino acid sequences were aligned, and then processed using Clustal Omega software [22], which calculated the percentage identity matrix. These percentage identity values were organized in a matrix format, facilitating the visualization of the relationships between the ANKH amino acid sequences across various species.

### 2.3 Zebrafish maintenance

Mature Tübingen (TU) zebrafish were raised in the Department of Physiology and Pharmacology, Western University, London, ON. The fish were kept in an aquarium at a temperature of 28°C, with a lighting schedule of 14 hours of light and 10 hours of darkness. The aquarium was supplied with fresh water and aeration to maintain its environment. The present study adhered to ethical guidelines established by the Canadian Council on Animal Care and was approved by the Animal Care Committee at Western University (AUP #2016-005).

### 2.4 mRNA Isolation, cDNA Synthesis, and Reverse Transcription RT-PCR

To determine the temporal expression pattern of *ankha* and *ankhb* we performed RT-PCR targeting 19 different developmental time points. Total mRNA was extracted from zebrafish including embryonic and larval developmental stages (2-cell (0.75 hours post fertilization (hpf)), 256 cell (2.5 hpf), sphere (4.0 hpf), dome (4.3 hpf), 50% epiboly (5.3 hpf), shield (6.0 hpf), 75% epiboly (8.0 hpf), 5-9 somites (11.0 hpf), 14-19 somites (16.0 hpf), 26+ somites (22.0 hpf), prim-5 (25.0 hpf), prim-15 (32.0 hpf), prim-25 (35 hpf), 2 dpf, 3 dpf, 4 dpf, 5 dpf, 6 dpf and to larval day 7. A pool of 30 embryos or larvae was collected for each stage, and mRNA extraction was performed using TRIzol reagent and chloroform (Invitrogen, 15596026). The isolated RNA was reverse transcribed into cDNA using the High-Capacity cDNA Reverse Transcription Kit, following the manufacturer’s instructions (Applied Biosystems, 4368814). Primer sets for *ankha* and *ankhb* were used for RT-PCR analysis, and *lsm12b* was used as a loading control [23] due to its relatively consistent expression levels throughout development (Table 1). Densitometry analysis was conducted using expression of each gene relative to the control (ImageJ).

**Table 1.**
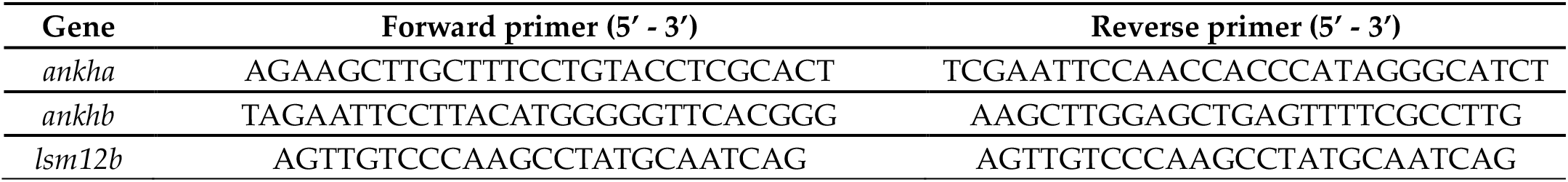
Primer sequences for zebrafish RT-PCR target genes.

### 2.5 Reverse Transcription - quantitative PCR (RT-qPCR)

RT-qPCR was conducted to quantify the gene expression patterns of *ankha* and *ankhb* on zebrafish samples at 2 dpf, 3 dpf, 4 dpf and 7 dpf, as these developmental stages represent times preceding, during and after bone formation. cDNA preparation was conducted as described above and primers and PCR conditions were selected in accordance with those previously described (Table 2) [24]. The comparative Ct method was used to analyze the relative gene expression, represented as 2^−(ΔΔ*Ct*)^, providing a ratio to the constitutively active *gapdh* expression.

**Table 2.**
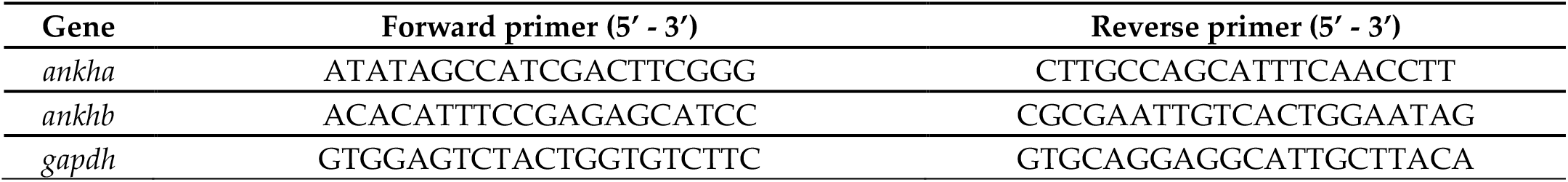
Primer sequences for zebrafish RT-qPCR target genes.

### 2.6 RNA Probe Generation for in situ Hybridization

Primers for *ankha* and *ankhb* were designed for *in situ* hybridization, and restriction enzyme sites were included for directional cloning. Riboprobes were synthesized from amplified *ankha* and *ankhb* fragments, digested using EcoRI and HindIII (New England Biolabs, R3101S and R0104S, respectively) and then ligated into the restriction enzyme-digested Bluescript SK+ plasmid. Following transformation into *E. coli* (DH5α), colonies were selected on LB/Agar plates containing ampicillin and subsequently identified using colony PCR. Finally, restriction enzyme digests were performed after plasmids were isolated using a midiprep kit (QIAGEN, 27104). All riboprobes were generated from linearized plasmids and synthesized using the DIG RNA Labelling Kit (Roche, 11175025910).

### 2.7 Whole-Mount in situ Hybridization (WISH)

*In situ* hybridization was performed to study the spatial expression patterns of *ankha* and *ankhb* genes in zebrafish larvae fixed at 2 dpf, 3 dpf and 4 dpf in 4% paraformaldehyde using an established protocol [25] Briefly, the fixed samples were incubated with a hybridization solution containing sense or antisense riboprobes. This step was followed by thorough washing, blocking with 5% heat-inactivated sheep serum and incubation with an anti-digoxigenin antibody conjugated to alkaline phosphatase (Invitrogen, AM1326). NBT/BCIP (4-nitro blue tetrazolium chloride/5-bromo-4-chloro-3-indolylphosphate, Roche, 11681451001) was added after thoroughly washing samples, and staining was carried out in the dark until a signal was detected in embryos/larvae incubated with sense strand probes. An antisense *krox-20* riboprobe (not shown) was used on zebrafish at 24 hpf as a positive control. All samples were cleared in glycerol and stored in PBS-Tween 20. Images were collected using a Nikon Inverted T12F Deconvolution Microscope (Biotron, Western University).

## 3. Results

### 3.1 The ankhb amino acid sequence in zebrafish demonstrates a closer evolutionary relationship to ANKH proteins of other vertebrates

Our first goal was to identify which of the zebrafish ankh proteins was more closely related to the single protein present in the “higher vertebrates.” The resulting neighbor-joining tree topology was aligned (Figure 1a), and the tree was rooted with the jawless fish species, hagfish (*Myxini)*. Within this tree, the Elephant shark (*Callorhinchus milii*), representing cartilaginous fish, branched off separately from the vertebrate group. Teleost species including zebrafish (*Danio rerio*), Mexican tetra (*Astyanax mexicanus*), and the common carp (*Cyprinus carpio*) formed a cohesive cluster with distinct monophyletic groups for ankha and ankhb. All other vertebrates are positioned on the phylogenetic tree in accordance with the established vertebrate phylogeny [26]. According to the neighbor-joining tree analysis the branch length of ankh from *Gallus gallus*, the common chicken, appeared to be evolving more rapidly compared to other vertebrate species.

To further elucidate the relationship between zebrafish ankha and ankhb and their counterparts in humans and mice, a maximum likelihood phylogenetic tree was constructed using these four sequences (Figure 1b). Zebrafish ankhb clustered within the same monophyletic group as the sequences from humans and mice, underscoring their close evolutionary relationship. In contrast, zebrafish ankha appeared more distantly related from higher vertebrates. The *per cent* identity values confirmed this observation (Figure 1c), revealing 83% amino acid sequence identity between zebrafish ankhb and human ANK and 81% identity to the mouse protein. In comparison, ankha had 77% and 76% sequence identity to that of human ANK and mouse Ank, respectively. These findings indicate that the ‘b’ paralog in zebrafish is evolutionarily closer to the ANK/Ank in humans and mice. To confirm that paralog ‘a’ was more ancient than ‘b’, we also analyzed the amino acid sequences of ankha and ankhb with sequences of ankh present in species that evolved before zebrafish. We did not identify invertebrate ankh from the Ensembl site, but ankh protein was identified in jawless fish species, specifically *Petromyzon marinus* and *Myxini*. Additionally, *Callorhinchus milii*, a cartilaginous fish, also harbors a single ankh protein, but evidence indicates that it would be characterized as ankhb. Hagfish exhibits the same level of identity (≈32%) for both ankha and ankhb, whereas *Petromyzon marinus* possessed only one *ankh* protein. Furthermore, this *P. marinus* transcript demonstrates a higher amino acid sequence identity to the zebrafish ankhb (Figure 1c). In summary, zebrafish ankhb demonstrates a closer evolutionary relationship to the ANK proteins of higher vertebrates and appears to represent the more ancient lineage among the ankh family.

### 3.2 ankha and ankhb Display Differential Expression Over Zebrafish Embronic and Larval Development

*RT-PCR* was performed to investigate the temporal expression of the *ankha* and *ankhb* genes with *lsm12b as a loading control* (Figure 2a). The expression of each paralog was seen in all embryonic and larval stages of development, and Sanger sequencing confirmed that they represent *ankha* and *<u>ankhb</u>*. Amplicons for both genes were detected at the 2-cell stage (0.75 hpf), showing that they are maternally derived and present before the midbastula transition [27]. *ankha* expression was higher than *ankhb* at the 2-cell stage, followed by a decrease in expression after 75% epiboly (8 hpf). An increase in *ankha* expression resumed at larval developmental stages (2 dpf, 3dpf, 4dpf, and 7 dpf) and was present at later stages. *ankhb* expression gradually increased from the 2-cell stage through to the larval stages. It should be noted that there was a decrease in *ankha* expression stage and a concomitant increase in *ankhb* expression at 75% epiboly. Despite this noticeable difference, amplicons corresponding to both *ankha* and *ankhb* gene were detected at all stages of development.

**Figure 2.**
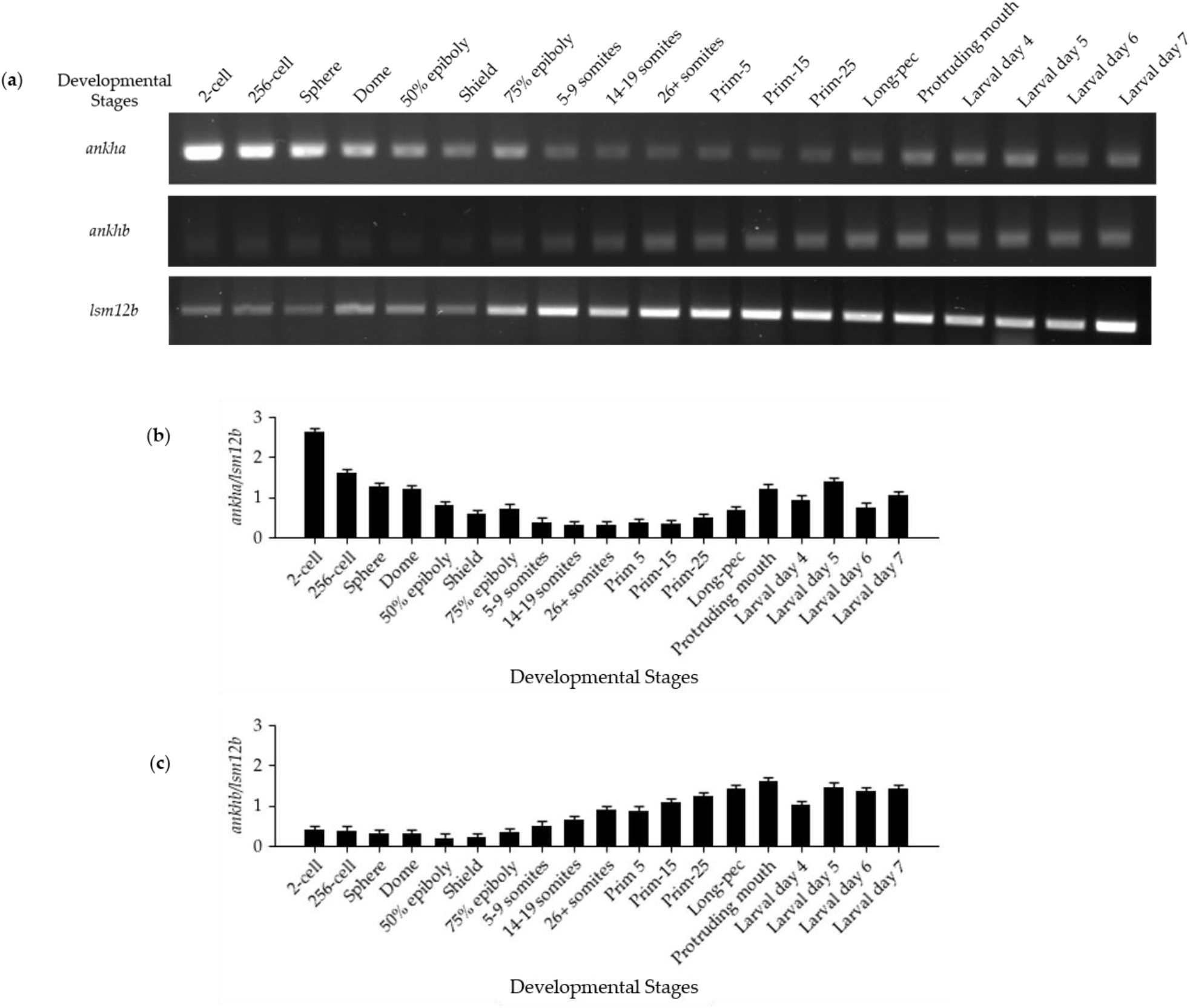
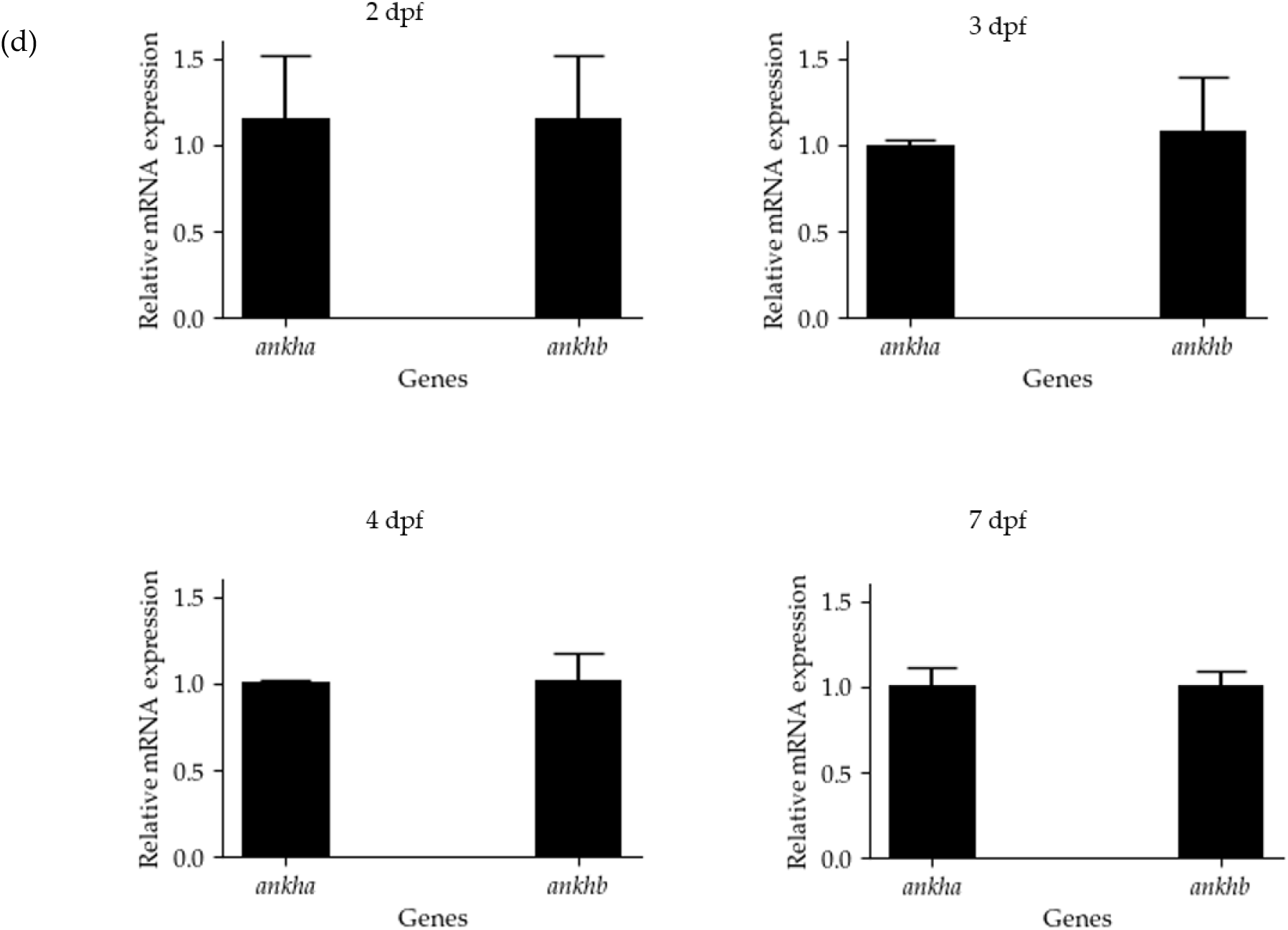
Temporal expression of *ankha* and *ankhb* during zebrafish development **(a)** Expression of *ankha* and *ankhb* relative to the *lsm12b* control in this representative endpoint PCR gel, N=3 replicates. Thirty embryos or fry from each developmental stage, ranging from the 2-cell stage (0.75 hpf) to 7 dpf, were sampled. **(b-c)** Semi-quantitative analysis of the temporal expression of *ankha* **(b)** and *ankhb* **(c)** in zebrafish, with each bar representing the expression of *ankha* and *ankhb* relative to the *lsm12b* expression. **(d)** RT-qPCR normalized expression of *ankha* and *ankhb* genes at 2, 3, 4 and 7 days post-fertilization (dpf) relative to *gapdh*, showing the mean +/- the standard error. Each sample represents 30 pooled larvae, where N=3 and statistical significance was determined using an unpaired t-test (p<0.05).

Densitometric analysis was used as a method to quantify gene expression observed in Figure 2a and to observed the differential gene expression pattern in *ankha* and *ankhb* during 19 different developmental stages (Figure 2b & c), there appeared to be different pattern in the expression of the *ankha* and *ankhb* genes. For a more quantified approach, RT-qPCR was used to detect gene expression during a temporal window that corresponds with bone formation. Specifically, we assessed *ankha* and *ankhb* expression at 2 dpf, a time just preceding bone formation, as well several stages during bone formation (3 dpf, 4 dpf, and 7 dpf). Expression of *ankhb* was higher than *ankha* expression throughout these stages, though this difference was not statistically significant. Given that *ankha* and *ankhb* are expressed throughout embryonic and larval development including stages that correspond to bone formation, we are unable to assign distinct developmental functions to each paralog.

### 3.3 Expression of ankha and ankhb is localized to the craniofacial region, somites, notochord, and tail

The spatial expression patterns of the *ankha* and *ankhb* genes during zebrafish larval development at 2 dpf, 3 dpf, and 4 dpf were examined using DIG-labeled antisense probes and WISH. Results at 2 dpf show staining of *ankha* and *ankhb* near the developing eye and notochord, with non-specific staining throughout the body. By day 3, both *ankha* and *ankhb* expression remained in the craniofacial region, specifically the brain, with somites and the notochord showing more localized staining compared to the remainder of the body. At 4 dpf, the expression pattern of *ankha* extended rostrally below the eyes and brain (star), with staining seen in the notochord and somites. The expression of *ankhb* at 4 dpf was slightly different for *ankha*, and staining was less intense in the brain and spinal cord and no signal seen below the eyes. Staining of *ankhb* was also present in the notochord and somites, similar to that seen for *ankha*.

## 4. Discussion

Human craniometaphyseal dysplasia (CMD) is a rare bone disorder caused by mutations in the *ANKH* gene. There is no permanent cure for this disorder; patients often require repeated recontouring of affected bones [4]. This and other CMD-associated problems highlight the significance of our research, and as a preliminary step we have taken advantage of the zebrafish attributes to model this rare disease. Our study focused on three pivotal areas: 1) the evolutionary lineage of the zebrafish *ankh* paralogous genes; 2) the similarities and differences in their temporal expression patterns; and 3) the spatial distribution of each transcript during development. In brief, we found a closer evolutionary relationship exists between the zebrafish ankhb protein and its human and other “higher” vertebrate counterparts. Furthermore, we noted distinct temporal expression patterns with *ankha* more prominently expressed in early development stages, and *ankhb* expression at larval growth stages. However, we were unable to assign the function of each paralog based on this analysis as there was significant overlap between the expressions of these genes. Hence, whole mount *in situ* hybridization was employed, hoping the spatial expression patterns of these paralogs would provide clues as to their function. Again, our analysis noted some differences in the spatial expression patterns of *ankha* and *ankhb*, especially in craniofacial regions. However, these patterns were subtle and provided no additional clues as to the functional involvement of each gene product.

To investigate the evolutionary conservation of ankha and ankhb to other vertebrates, our analysis spanned multiple vertebrate phyla, with a specific focus on comparisons between human and mouse variants. Two paralogous *ankh* genes were identified in zebrafish and they arose from a genome-wide duplication event (Fig. 1a). Initially, there were two significant whole-genome duplication events, which occurred during vertebrate evolution [28]. A third event occurred and this was in teleost, specifically involving a subgroup of the ray-finned fishes (Actinopterygii) [29]. Nevertheless, our comparisons revealed that ankhb showed a closer evolutionary relationship to the ANK/Ank present in higher vertebrates (Fig.1a). Furthermore, ankha had diverged more, as indicated by the branch lengths in the neighbor-joining tree (Fig 1a). This evolutionary pattern highlights the role of gene duplication in facilitating the emergence of novel protein function [30], but there are other cases where duplication results in the same function of the protein,. For example, zebrafish insulin-like growth factor (*IGF*) genes, those identified as *igf-1a, igf-1b, igf-2a*, and *igf-2b*, which produce four structurally and functionally unique IGF peptides. Humans typically possess only one IGF-1 and one IGF-2 gene, which primarily regulate growth and development. Functional studies reveal that all four *igf*s induce similar developmental abnormalities, with varying degrees of severity, including full or partial duplication of notochords. These findings shed light on the functional similarities among the duplicated *igf* genes [31]. However, over time this redundancy enables one of the paralogs to evolve with less selective pressure and this freedom allows for the accumulation of mutations [32]. While such mutations might be harmful to a gene that exists in only one copy, they can sometimes result in the emergence of new functions for the duplicated gene [33] A severe form of childhood epilepsy results from mutations in the *SCN1A* gene in humans. In Zebrafish, two paralog genes which are homologous to human *SCN1A*: *scn1laa* and *scn1lab* have been identified. Disrupting one of these paralogs in zebrafish mimics a heterozygous state of the disease in humans, as the other paralog is still present. Despite the widespread acceptance of the ‘disease-state model,’ evidence indicates that the functionality of these genes might not be entirely similar. Several non-conserved hotspots were identified within the scn1aa and scn1ab proteins, which lead to functional differences. This further validates that the accumulation of mutations in paralog genes can result in distinct functions [34]. The observed divergence between ankha and ankhb may reflect such an evolutionary process, where differential selective pressures and mutational landscapes have resulted in distinct evolutionary trajectories and functional specializations within vertebrate lineages [33]. We found that the evolutionary history of the ankh gene family reveals intriguing patterns of conservation and divergence across vertebrate lineages (Fig.1a). Our analysis would indicate that a single ankh protein was present in species predating some teleost fish as well as in higher vertebrates. Although this protein is closely related to zebrafish ankhb, it suggests that the ankhb paralog may represent a foundational genetic element retained from ancestral vertebrates [20], potentially preserved through the first or second round of whole-genome duplication. This persistence of the ankhb paralog highlights its evolutionary importance, and suggests a selective advantage that favored its conservation through significant evolutionary transitions.

Having determined that zebrafish ankhb was more closely related to the ANK/Ank in human and mouse, our analysis focused on the temporal expression patterns of zebrafish *ankha* and *ankhb* (Fig. 2a). If one paralog was expressed before the other, it might suggest differing functions. Despite this, the human and mouse *ANK*/*Ank* gene encodes a single protein also involved in ATP and other metabolite transport occurring in human embryonic kidney cells [7]. Our study employed 19 distinct developmental time points, aimed to determine when *ankha* and *ankhb* were expressed during zebrafish development. Furthermore, since our focus was on bone mineralization and eventually modeling human CMD, we used zebrafish stages that preceded and followed bone development. The temporal expression patterns for *ankha* and *ankhb* revealed both genes were present from the 2-cell stage, corroborating previous studies [10, 19] and highlighting the maternal inheritance of these genes. Subtle differences were noted (Fig. 2a), however, with *ankha* expression being higher than *ankhb* at early stages, prior to bone mineralization, but we can only speculate on their roles in development as there was considerable overlap in each pattern. The presence of *ankha* and *ankhb* transcripts at stages before bone mineralization (Fig. 2a, 2b and 2c), would suggest that both genes are implicated in additional functions. In fact, human oocytes exhibit the highest levels of ANK among different cell types, and high expression has also been noted in oligodendrocytes and inhibitory neurons [35]. These maternal gene products regulate various processes that eventually leads to the activation of the zygotic genome [36]. While it is not known whether zebrafish *ankh* genes regulate these critical processes before the midblastula transition, our research would indicate they are present in brain tissue (Fig. 3). Furthermore, humans, mice, and pigs show ANK/Ank protein level patterns that are predominantly found in the brain, specifically in the cerebellum [35]. Brain tissue has high ATP demands [37] and zebrafish also contain a pool of maternally inherited ATP [38, 39]. If *ankha* and *ankhb* encode proteins required for ATP transport, it might explain why the gene transcripts are present at these early stages before zygotic activation at cycle 10 [27]. Studies also indicate that the period from the 3/4 and 4/5 hpf to 24 hpf is associated with a substantial rise in metabolites, including amino acids and nucleic acids, and a significant increase in energy metabolites like complex sugars and TCA cycle intermediates [40]. These intermediates would also contribute to the increased levels of ATP and other metabolites and the presence of *ankha* and *ankhb* transcripts, if translated, would likely be involved in their transport. Whether both genes in zebrafish contribute to transporting ATP and metabolites, or if they have diverged functionally, remains to be determined.

**Figure 3.**
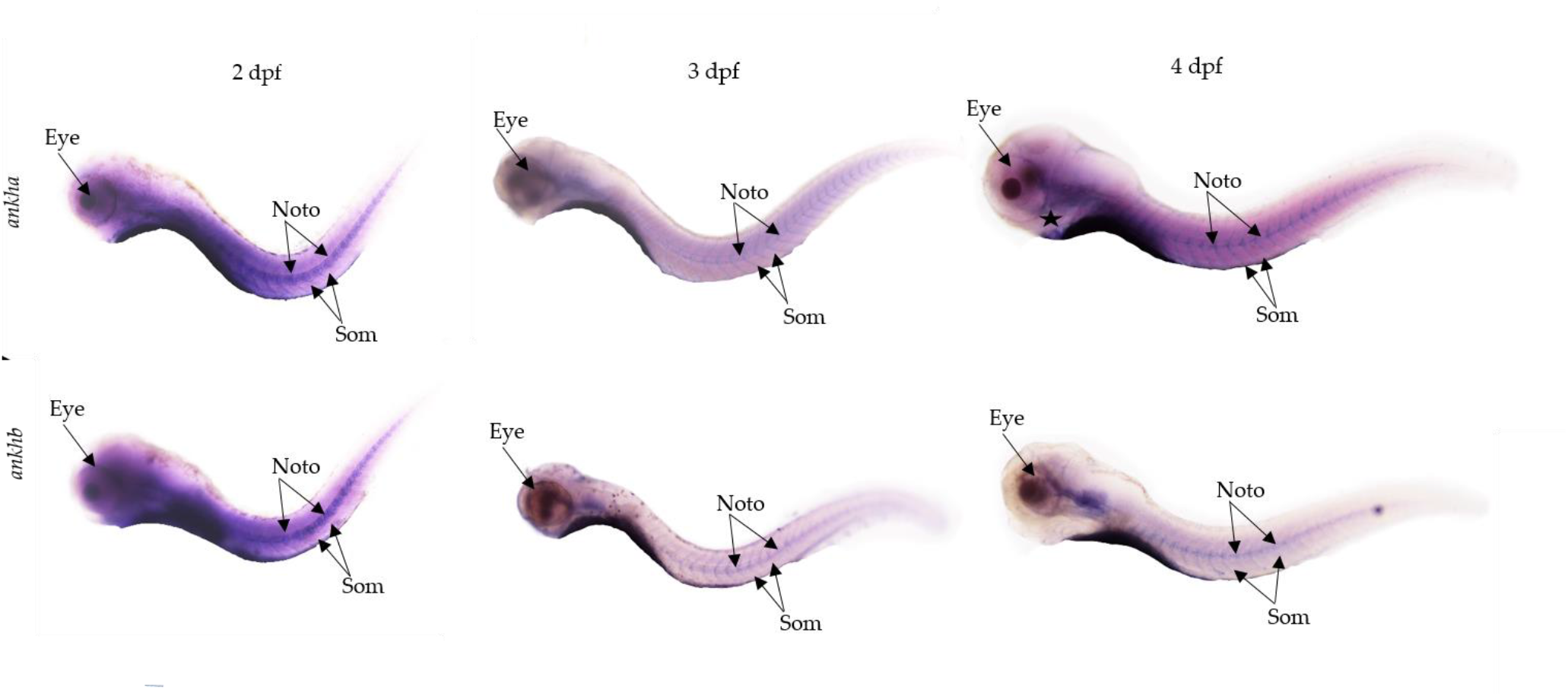
Whole mount *in situ* hybridization analysis of *ankha* and *ankhb* spatial expression patterns in zebrafish development. Lateral views of zebrafish larval stages at 2, 3, and 4 days post-fertilization (dpf) reveal spatial gene expression patterns for *ankha* and *ankhb*. At 2 and 3 dpf, both *ankha* and *ankhb* show predominant staining in the craniofacial region, somites and notochord. At 4 dpf, *ankha* extends below the eye (star), and appears in the somites, and notochord (arrows). *ankhb* expression at 3 dpf and 4 dpf is mainly localized to the craniofacial region and notochord area. For *ankhb*, staining below the eye is reduced and is more localized in regions below the hindbrain. Noto: Notochord, Som: Somites. Scale bar indicate 100 um.

ANK/Ank is known to control of pyrophosphate levels [8] and *ankha* and *ankhb* are expressed at the time when bone mineralization occurs in zebrafish [14]. Our results revealed *ankha* and *ankhb* were present in the zebrafish craniofacial region (Fig. 3), particularly below the brain where intramembranous and endochondral bone will develop [16]. Since ANK/Ank is known to regulate bone mineralization [8], it is imperative to determine the role of each paralog gene in zebrafish. If one or the other zebrafish paralog, or both, is/are involved in bone mineralization then CRISPR-Cas9 editing tools and other *in vitro* methods can be employed to assign roles to these paralogs. However, at present, *ankha* and *ankhb* show overlapping spatiotemporal expression patterns and their individual functions remain unknown.

## 5. Conclusion

Our research provides significant insights into the zebrafish *ankha* and *ankhb* genes, underscoring their potential implications for understanding and addressing craniometaphyseal dysplasia (CMD) in humans. We discovered a profound evolutionary link between the zebrafish ankhb amino acids and those in higher vertebrates, demonstrating the evolutionary persistence of this gene. Our data also showed that *ankha* and *ankhb* are maternally inherited and that the expression of *each gene* occurs during the onset of larval development and bone mineralization. Notably, the high expression of these genes in early developmental stages and their spatial localization to the craniofacial region suggest a role in ATP transport, which is crucial due to the high ATP demand in the craniofacial region with development of the brain. However, without detailed functional analyses, the exact roles of these genes remain speculative. The subsequent generation of *ankha* and *ankhb* knockouts using CRISPR-Cas9 gene editing methods will elucidate their roles in development and provide a platform to address how we can manage CMD in the human population. This study lays the groundwork for future research into the genetic mechanisms of bone diseases and the potential for targeted therapies in CMD.

## Acknowledgments

The authors would like to acknowledge Dr. D.R. Smith (Biology, Western University) for discussions and assistance with the phylogenetic analysis.

## References

[1] S. R. Kim and Y. S. Han, “Craniometaphyseal Dysplasia,” Arch Plast Surg, vol. 40, no. 2, pp. 157–159, Mar. 2013, doi: 10.5999/aps.2013.40.2.157.

[2] J. Wu, X. Li, and S. Chen, “Special manifestations and treatment of rare cases of snoring with special facial features and hearing loss in children,” J Int Med Res, vol. 50, no. 7, p. 03000605221108085, Jul. 2022, doi: 10.1177/03000605221108085.

[3] E. Reichenberger et al., “Autosomal Dominant Craniometaphyseal Dysplasia Is Caused by Mutations in the Transmembrane Protein ANK,” The American Journal of Human Genetics, vol. 68, no. 6, pp. 1321–1326, Jun. 2001, doi: 10.1086/320612.

[4] I.-P. Chen, C. J. Wang, S. Strecker, B. Koczon-Jaremko, A. Boskey, and E. J. Reichenberger, “Introduction of a Phe377del Mutation in ANK Creates a Mouse Model for Craniometaphyseal Dysplasia,” Journal of Bone and Mineral Research, vol. 24, no. 7, pp. 1206–1215, Jul. 2009, doi: 10.1359/jbmr.090218.

[5] J. P. Stains and R. Civitelli, “Connexins in The Skeleton,” Semin Cell Dev Biol, vol. 50, pp. 31–39, Feb. 2016, doi: 10.1016/j.semcdb.2015.12.017.

[6] P. Nürnberg et al., “Heterozygous mutations in ANKH, the human ortholog of the mouse progressive ankylosis gene, result in craniometaphyseal dysplasia,” Nat Genet, vol. 28, no. 1, pp. 37–41, May 2001, doi: 10.1038/ng0501-37.

[7] F. Szeri et al., “The membrane protein ANKH is crucial for bone mechanical performance by mediating cellular export of citrate and ATP,” PLoS Genet, vol. 16, no. 7, p. e1008884, Jul. 2020, doi: 10.1371/journal.pgen.1008884.

[8] F. Szeri et al., “The Mineralization Regulator ANKH Mediates Cellular Efflux of ATP, Not Pyrophosphate,” J Bone Miner Res, vol. 37, no. 5, pp. 1024–1031, May 2022, doi: 10.1002/jbmr.4528.

[9] R. A. Terkeltaub, “Inorganic pyrophosphate generation and disposition in pathophysiology,” American Journal of Physiology-Cell Physiology, vol. 281, no. 1, pp. C1–C11, Jul. 2001, doi: 10.1152/ajpcell.2001.281.1.C1.

[10] Y.-J. Ding, X.-G. Chen, and Y.-H. Chen, “Molecular structure and developmental expression of two zebrafish Ankylosis Progressive Homolog (ankh) genes, ankha and ankhb,” Russ J Dev Biol, vol. 44, no. 6, pp. 307–313, Nov. 2013, doi: 10.1134/S1062360413060106.

[11] I.-P. Chen, L. Wang, X. Jiang, H. L. Aguila, and E. J. Reichenberger, “A Phe377del mutation in ANK leads to impaired osteoblastogenesis and osteoclastogenesis in a mouse model for craniometaphyseal dysplasia (CMD),” Hum Mol Genet, vol. 20, no. 5, pp. 948–961, Mar. 2011, doi: 10.1093/hmg/ddq541.

[12] A. M. Ho, M. D. Johnson, and D. M. Kingsley, “Role of the Mouse ank Gene in Control of Tissue Calcification and Arthritis,” Science, vol. 289, no. 5477, pp. 265–270, Jul. 2000, doi: 10.1126/science.289.5477.265.

[13] K. A. Gurley, R. J. Reimer, and D. M. Kingsley, “Biochemical and Genetic Analysis of ANK in Arthritis and Bone Disease,” Am J Hum Genet, vol. 79, no. 6, pp. 1017–1029, Dec. 2006.

[14] F. Tonelli et al., “Zebrafish: A Resourceful Vertebrate Model to Investigate Skeletal Disorders,” Frontiers in Endocrinology, vol. 11, 2020, Accessed: Sep. 11, 2023. [Online]. Available: https://www.frontiersin.org/articles/10.3389/fendo.2020.00489

[15] M. Carnovali, G. Banfi, and M. Mariotti, “Zebrafish Models of Human Skeletal Disorders: Embryo and Adult Swimming Together,” BioMed Research International, vol. 2019, p. e1253710, Nov. 2019, doi: 10.1155/2019/1253710.

[16] P. Le Pabic, D. B. Dranow, D. J. Hoyle, and T. F. Schilling, “Zebrafish endochondral growth zones as they relate to human bone size, shape and disease,” Front Endocrinol (Lausanne), vol. 13, p. 1060187, 2022, doi: 10.3389/fendo.2022.1060187.

[17] K. Dietrich et al., “Skeletal Biology and Disease Modeling in Zebrafish,” Journal of Bone and Mineral Research, vol. 36, no. 3, pp. 436–458, 2021, doi: 10.1002/jbmr.4256.

[18] S. G. Vascotto, Y. Beckham, and G. M. Kelly, “The zebrafish’s swim to fame as an experimental model in biology,” Biochem. Cell Biol., vol. 75, no. 5, pp. 479–485, Oct. 1997, doi: 10.1139/o97-081.

[19] R. J. White et al., “A high-resolution mRNA expression time course of embryonic development in zebrafish,” eLife, vol. 6, p. e30860, Nov. 2017, doi: 10.7554/eLife.30860.

[20] F. J. Martin et al., “Ensembl 2023,” Nucleic Acids Research, vol. 51, no. D1, pp. D933–D941, Jan. 2023, doi: 10.1093/nar/gkac958.

[21] K. Tamura, D. Peterson, N. Peterson, G. Stecher, M. Nei, and S. Kumar, “MEGA5: Molecular Evolutionary Genetics Analysis Using Maximum Likelihood, Evolutionary Distance, and Maximum Parsimony Methods,” Molecular Biology and Evolution, vol. 28, no. 10, pp. 2731–2739, Oct. 2011, doi: 10.1093/molbev/msr121.

[22] F. Sievers et al., “Fast, scalable generation of high-quality protein multiple sequence alignments using Clustal Omega.,” Mol Syst Biol, vol. 7, p. 539, Oct. 2011, doi: 10.1038/msb.2011.75.

[23] Y. Hu, S. Xie, and J. Yao, “Identification of Novel Reference Genes Suitable for qRT-PCR Normalization with Respect to the Zebrafish Developmental Stage,” PLoS One, vol. 11, no. 2, p. e0149277, Feb. 2016, doi: 10.1371/journal.pone.0149277.

[24] M. Van Gils, A. Willaert, E. Y. G. De Vilder, P. J. Coucke, and O. M. Vanakker, “Generation and Validation of a Complete Knockout Model of abcc6a in Zebrafish,” Journal of Investigative Dermatology, vol. 138, no. 11, pp. 2333–2342, Nov. 2018, doi: 10.1016/j.jid.2018.06.183.

[25] M. N. Knowlton, B. M. C. Chan, and G. M. Kelly, “The zebrafish band 4.1 member Mir is involved in cell movements associated with gastrulation,” Developmental Biology, vol. 264, no. 2, pp. 407–429, Dec. 2003, doi: 10.1016/j.ydbio.2003.09.001.

[26] J. R. Peat, O. Ortega-Recalde, O. Kardailsky, and T. A. Hore, “The elephant shark methylome reveals conservation of epigenetic regulation across jawed vertebrates.” F1000Research, Apr. 20, 2017. doi: 10.12688/f1000research.11281.1.

[27] D. A. Kane and C. B. Kimmel, “The zebrafish midblastula transition,” Development, vol. 119, no. 2, pp. 447–456, Oct. 1993, doi: 10.1242/dev.119.2.447.

[28] D. Steinke, S. Hoegg, H. Brinkmann, and A. Meyer, “Three rounds (1R/2R/3R) of genome duplications and the evolution of the glycolytic pathway in vertebrates,” BMC Biology, vol. 4, no. 1, p. 16, Jun. 2006, doi: 10.1186/1741-7007-4-16.

[29] E. M. Kollitz, M. B. Hawkins, G. K. Whitfield, and S. W. Kullman, “Functional Diversification of Vitamin D Receptor Paralogs in Teleost Fish After a Whole Genome Duplication Event,” Endocrinology, vol. 155, no. 12, pp. 4641–4654, Dec. 2014, doi: 10.1210/en.2014-1505.

[30] A. L. Hughes, “Gene duplication and the origin of novel proteins,” Proceedings of the National Academy of Sciences, vol. 102, no. 25, pp. 8791–8792, Jun. 2005, doi: 10.1073/pnas.0503922102.

[31] S. Zou, H. Kamei, Z. Modi, and C. Duan, “Zebrafish IGF Genes: Gene Duplication, Conservation and Divergence, and Novel Roles in Midline and Notochord Development,” PLOS ONE, vol. 4, no. 9, p. e7026, Sep. 2009, doi: 10.1371/journal.pone.0007026.

[32] S. P. Otto and P. Yong, “16 - The Evolution of Gene Duplicates,” in Advances in Genetics, vol. 46, J. C. Dunlap and C. -ting Wu, Eds., in Homology Effects, vol. 46., Academic Press, 2002, pp. 451–483. doi: 10.1016/S0065-2660(02)46017-8.

[33] F. A. Kondrashov, I. B. Rogozin, Y. I. Wolf, and E. V. Koonin, “Selection in the evolution of gene duplications,” Genome Biology, vol. 3, no. 2, p. research0008.1, Jan. 2002, doi: 10.1186/gb-2002-3-2-research0008.

[34] W. J. Weuring, J. W. Hoekman, K. P. J. Braun, and B. P. C. Koeleman, “Genetic and Functional Differences between Duplicated Zebrafish Genes for Human SCN1A,” Cells, vol. 11, no. 3, Art. no. 3, Jan. 2022, doi: 10.3390/cells11030454.

[35] M. Uhlén et al., “Tissue-based map of the human proteome,” Science, vol. 347, no. 6220, p. 1260419, Jan. 2015, doi: 10.1126/science.1260419.

[36] F. L. Marlow, “Introduction,” in Maternal Control of Development in Vertebrates: My Mother Made Me Do It!, Morgan & Claypool Life Sciences, 2010. Accessed: Nov. 06, 2023. [Online]. Available: https://www.ncbi.nlm.nih.gov/books/NBK53192/

[37] A. Faria-Pereira and V. A. Morais, “Synapses: The Brain’s Energy-Demanding Sites,” Int J Mol Sci, vol. 23, no. 7, p. 3627, Mar. 2022, doi: 10.3390/ijms23073627.

[38] A. Dutta and D. K. Sinha, “Zebrafish lipid droplets regulate embryonic ATP homeostasis to power early development,” Open Biol, vol. 7, no. 7, p. 170063, Jul. 2017, doi: 10.1098/rsob.170063.

[39] D. Calzia et al., “Evidence of Oxidative Phosphorylation in Zebrafish Photoreceptor Outer Segments at Different Larval Stages,” J Histochem Cytochem, vol. 66, no. 7, pp. 497–509, Jul. 2018, doi: 10.1369/0022155418762389.

[40] S. S. Dhillon et al., “Metabolic profiling of zebrafish embryo development from blastula period to early larval stages,” PLOS ONE, vol. 14, no. 5, p. e0213661, May 2019, doi: 10.1371/journal.pone.0213661.

